# Candidate Explorer: a tool for discovery, evaluation, and display of mutations causing significant immune phenotypes

**DOI:** 10.1101/2020.11.07.371914

**Authors:** Darui Xu, Stephen Lyon, Chun Hui Bu, Sara Hildebrand, Jin Huk Choi, Xue Zhong, Aijie Liu, Emre E. Turer, Zhao Zhang, Evan Nair-Gill, Hexin Shi, Ying Wang, Duanwu Zhang, Tao Yue, Jeff SoRelle, Takuma Misawa, Lei Sun, Jianhui Wang, Roxana Farokhnia, Andrew Sakla, Sara Schneider, Nathan Stewart, Hannah Coco, Gabrielle Coolbaugh, Braden Hayse, Sara Mazal, Dawson Medler, Brandon Nguyen, Edward Rodriguez, Andrew Wadley, Miao Tang, Xiaohong Li, Priscilla Anderton, Katie Keller, Lindsay Scott, Jiexia Quan, Sydney Cooper, Baifang Qin, Jennifer Cardin, Rochelle Simpson, Meron Tadesse, Qihua Sun, John Santoyo, Amy Bronikowski, Alexyss Johnson, Eva Marie Y. Moresco, Bruce Beutler

**Affiliations:** Center for the Genetics of Host Defense, University of Texas Southwestern Medical Center, Dallas, Texas 75390; Department of Immunology, University of Texas Southwestern Medical Center, Dallas, Texas 75390; Department of Internal Medicine, Division of Gastroenterology, University of Texas Southwestern Medical Center, Dallas, TX 75390

## Abstract

When applied to immunity, forward genetic studies use meiotic mapping to provide strong statistical evidence that a particular mutation is causative of a particular immune phenotype. Notwithstanding this, co-segregation of multiple mutations, occasional unawareness of mutations, and paucity of homozygotes may lead to erroneous declarations of cause and effect. We sought to improve the selection of authentic causative mutations using a machine learning software tool, Candidate Explorer (CE), which integrates 65 data features into a single numeric score, mathematically convertible to the likelihood of verification of any putative mutation-phenotype association. CE has identified most genes within which mutations can be causative of flow cytometric phenovariation in *Mus musculus*. The majority of these genes were not previously known to support immune function or homeostasis. Mouse geneticists will find CE data informative in identifying causative mutations within quantitative trait loci, while clinical geneticists may use CE to help connect causative variants with rare heritable diseases of immunity, even in the absence of linkage information. CE displays integrated mutation, phenotype, and linkage data, and is freely available for query online.

---

Forward genetics always begins with a phenotype, often induced by a random germline mutagen, and ends with the discovery of a causative mutation. We developed a process for rapid identification of causative mutations in mice carrying N-ethyl-N-nitrosourea (ENU)-induced germline mutations^1, 2^. Our pipeline involves mutagenizing male C57BL/6J (G0) mice and breeding them on the C57BL/6J background to create G1 male pedigree founders, G2 daughters, and G3 mice of both sexes for phenotypic screening (Supplementary Fig. 1). All G1 founders of pedigrees are whole-exome sequenced to identify >99% of ENU-induced non-synonymous coding/splicing changes, and all G2 and G3 mice are genotyped at these mutation sites in advance of phenotypic screening. Automated meiotic mapping (AMM) is performed using the program Linkage Analyzer, which tests the null hypothesis for every mutation in every screen; i.e., “mutation A is unrelated to phenotypic performance in screen α”^1^. Rejection of the null hypothesis with a *P* value ≤ 0.05, with Bonferroni correction for multiple comparisons, has generally been considered suggestive of causation. Verification by an independently generated allele is necessary to confirm the association.

Experience with many thousands of mutation-phenotype associations identified by AMM and either verified or excluded by testing CRISPR/Cas9-targeted alleles, has shown that the *P* value determined by AMM is not the sole indicator of causation. Many other factors, such as the nature of the mutation (benign, damaging, null), the essentiality of the gene for survival prior to weaning, pedigree size, the number of homozygotes tested, the magnitude of phenotypic effect, data variance characteristic of the screen in question, the number of distinct phenotypes caused by the mutation, the presence or absence of co-segregating mutations, and the observation of other alleles with similar effects, influence correct selection of an authentic causative mutation. These numerous considerations, not readily integrated into a decision by human observers, impelled us to develop Candidate Explorer (CE), a software tool employing a supervised machine learning algorithm to estimate the likelihood of verification of any putative mutation-phenotype association implicated by AMM.

In this study, we focused on changes in immune cell populations caused by ENU-induced mutations and detected by flow cytometric analysis of peripheral blood leukocytes from G3 mutant mice. We present CE assessments of 81,760 mutation-phenotype associations (*P*< 0.05). CE has identified more than 1,000 genes with a high and defined probability of verifiable importance in leukocyte development or maintenance. Many of these genes were not previously known to be important in immune function.

## Results

### CE overview

The purpose of CE is to aid the researcher in predicting whether a mutation associated with a phenotype by AMM is a truly causative mutation. CE evaluates mutation-phenotype associations that pass specific basal filters for conventionally good candidates. In this paper, we use as the default filters: *P*<0.05 (Bonferroni corrected), ≥ 10 mice in the tested pedigree, and ≥ 2 homozygous reference mice screened; however, more stringent criteria can be set by the user. The core of CE is a supervised machine learning algorithm that outputs a numerical score (ML Score) and a categorical assessment (Candidate Status) of each mutation-phenotype association based on input phenotype data (from screening), mutation data, gene data, and meiotic mapping data (Fig. 1a). CE is trained based on phenotypic assessment of mice carrying targeted null or replacement alleles of candidate genes (see below). In training, performed four times per day because of the dynamic status of the database, CE associates all defined features of the original pedigree screening data with positive or negative outcomes in the assessment of phenotypes in a pedigree of gene-targeted mice.

**Figure 1.**
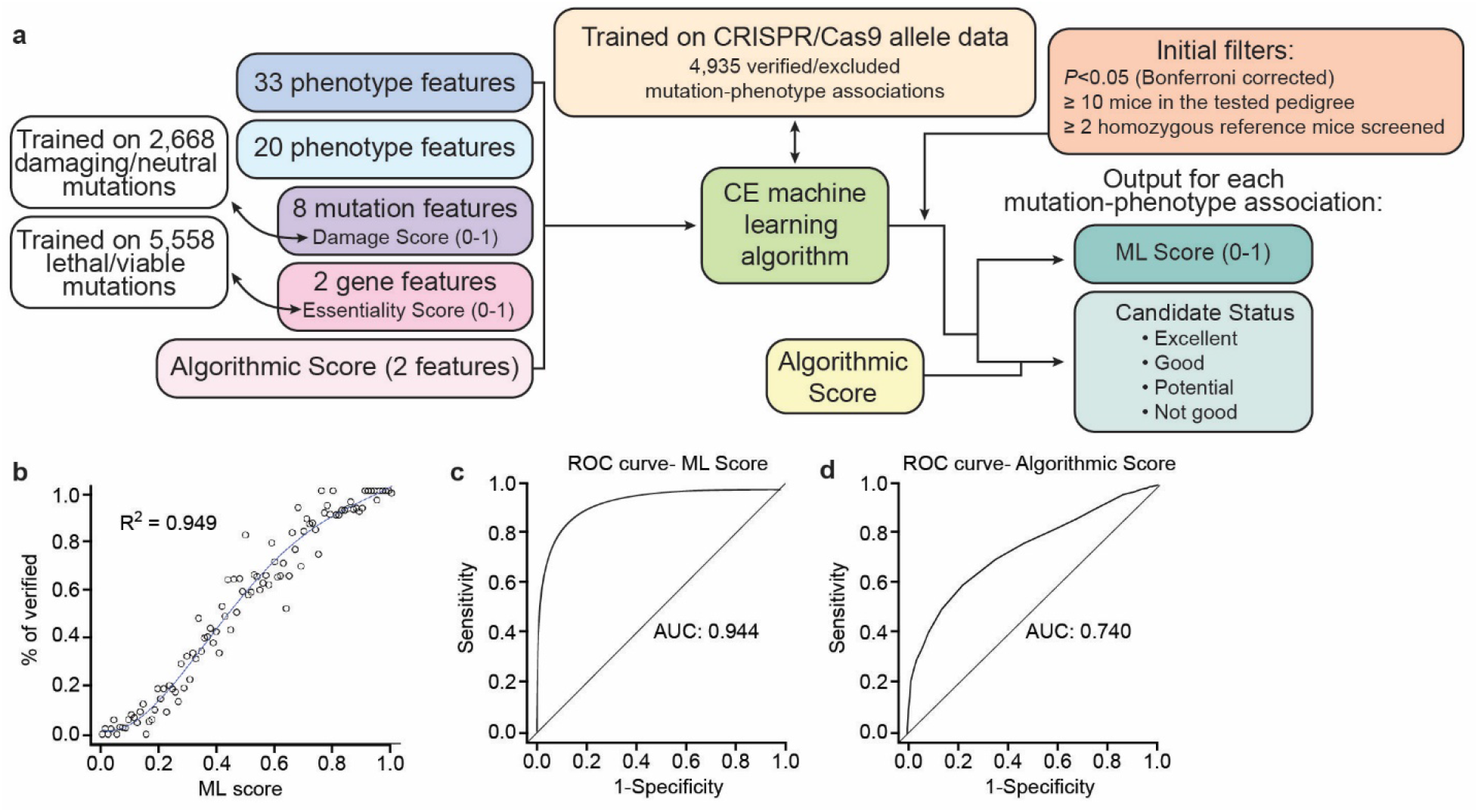
Candidate Explorer overview and performance. (**a**) Schematic of input features and outputs from CE. (**b**) Polynomial regression analysis of ML Score and average percentage of verified mutation-phenotype associations. N=4,893 mutation-phenotype associations and 487 CRISPR/Cas9-targeted genes. (**c,d**) ROC curves for ML Score (**c**) and Algorithmic Score (**d**).

CE is publicly available for querying mutation-phenotype associations identified in flow cytometry screens we have performed to date. An example of the use of CE is presented in Supplementary Video 1.

### Training and performance of CE

At present, the CE training set contains 1,990 verified and 2,945 excluded mutation-phenotype associations (4,935 assessments in all), based on germline retargeting of 490 genes. Germline retargeting was performed using CRISPR/Cas9 to generate knockout allele(s) of the candidate genes in mice on a pure reference background (C57BL/6J or C57BL/6N). Alternatively, when evidence for homozygous lethality of null alleles existed (see Essentiality Score) or the ENU mutation was suspected to cause hypermorphic, neomorphic, or antimorphic effects, the original ENU allele, typically a point mutation, was re-created by CRISPR/Cas9 targeting (designated “replacement” allele). Mice carrying targeted germline knockout or replacement alleles were expanded to form pedigrees containing mice homozygous for the reference allele (REF), heterozygous (HET), and homozygous for the variant allele (VAR). Compound heterozygous mice with two variant alleles of a gene were sometimes also generated. Fresh pedigrees of mice carrying the CRISPR-targeted alleles were subjected to the phenotypic screens in which the original ENU mutations scored as hits. CRISPR-targeted mutations were considered verified according to the criteria:

1. Observation of the same phenotype with the same directionality of change as observed for the original ENU allele with a P value better than 0.01, or
2. Observation of the same phenotype with the opposite directionality of change as observed for the original ENU allele with a P value better than 0.001, or
3. *De novo* observation of a phenotype (not seen in the original screen) with a P value better than 0.001.

The machine learning (ML) Score (range 0-1) output by CE is a class probability related by a polynomial function to the actual probability of verification by CRISPR-targeted alleles, as determined by regression analysis (Fig. 1b). In conjunction with the Algorithmic Score, it is used by CE to designate one of four possible Candidate Statuses for each mutation-phenotype association (excellent, good, potential, or not good). We generally choose good or excellent candidates for CRISPR/Cas9 targeting and further study. However, ML Scores are not strictly proportional to the probability of verification (Fig. 1b) and some “good” or “excellent” candidates fail to verify. Conversely, “potential” and “not good” candidates will sometimes verify as true positive associations. We take it as a truism that authentic candidates will achieve strong ML Scores as more alleles are obtained and tested (approaching saturation) and will therefore eventually be verified.

The performance of the CE prediction model established using the training set was assessed using the repeated 10-fold cross-validation method. The receiver operating characteristic (ROC) curve has an area under the curve (AUC) of 0.944 (Fig. 1c); the current cutoff is 0.43, corresponding to the point with the minimum distance to the upper left corner of the ROC curve. CE ranking of “good” or better corresponds to approximately 83% precision (correctly calling a verified candidate “true;” i.e., a 17% false discovery rate) and 85% recall (true positive rate) (Table 1).

**Table 1.** CE Performance

| **Mutation-phenotype associations (N=82,373; 1,050 verified, 1,393 excluded, 79,930 untested)** |  |  |  |  |  |  |  |
| --- | --- | --- | --- | --- | --- | --- | --- |
| **Cutoff** | **True positives** | **False positives** | **True negatives** | **False negatives** | **Recall** | **Accuracy** | **Precision** |
| Excellent | 656 | 107 | 1286 | 394 | 62.48% | 79.49% | 85.98% |
| Good and above | 899 | 183 | 1210 | 151 | 85.62% | 86.33% | 83.09% |
| Potential and above | 967 | 509 | 884 | 83 | 92.10% | 75.77% | 65.51% |
| Not good and above | 1050 | 1393 | 0 | 0 | 100% | 42.98% | 42.98% |
| **Alleles (N=2,252 named alleles; 225 verified, 84 excluded, 1,943 untested)** |  |  |  |  |  |  |  |
| Excellent | 167 | 10 | 74 | 58 | 74.22% | 77.99% | 94.35% |
| Good and above | 197 | 16 | 68 | 28 | 87.56% | 85.76% | 92.49% |
| Potential and above | 208 | 48 | 36 | 17 | 92.44% | 78.96% | 81.25% |
| Not good and above | 225 | 84 | 0 | 0 | 100% | 72.82% | 72.82% |
| **Genes (N=14,437; 126 causative, 89 non-causative, 14,222 untested)** |  |  |  |  |  |  |  |
| Excellent | 91 | 9 | 80 | 35 | 72.22% | 79.53% | 91.00% |
| Good and above | 112 | 14 | 75 | 14 | 88.89% | 86.98% | 88.89% |
| Potential and above | 120 | 52 | 37 | 6 | 95.24% | 73.02% | 69.77% |
| Not good and above | 126 | 89 | 0 | 0 | 100% | 58.60% | 58.60% |

CE is also often capable of correctly identifying which mutation is causative when two or more mutations co-localize (see also below, Driven By status). Among 969 such cases, CE correctly identified on average 87.3% of causative mutations as the top ML scorer, with generally better performance when fewer mutations colocalized (Table 2). As further training is performed, and as the total volume of screening data increases (with an attendant increase in the number of genes with allelism and the overall density of allelic series), CE performance will continue to improve.

**Table 2.** CE performance in scoring colocalizing mutation

| **# of co-localized genes** | **Total cases** | **CE rank correctly** | **% Correct** |
| --- | --- | --- | --- |
| 2 | 595 | 520 | 87.5% |
| 3 | 206 | 210 | 90.5% |
| 4 | 88 | 81 | 85.3% |
| 5 | 35 | 25 | 71.4% |
| 6 | 5 | 4 | 80.0% |
| 7 | 6 | 5 | 83.3% |
| 8 | 1 | 1 | 100.0% |
| 10 | 1 | 1 | 100.0% |

### Input data features

The CE prediction model was built using a random forest algorithm implemented in the R package and currently incorporates 65 features of the input data (33 phenotype features, 20 linkage analysis features, 8 mutation features, 2 gene features, and 2 other features; Table 3). The 20 most important features are ranked in Table 4. The Damage Score and Essentiality Score result from independent machine learning programs. The rule-based Algorithmic Score results from computational execution of a fixed algorithm that was human designed.

**Table 3.** Features of input data in CE prediction algorithm

|  |
| --- |
| **Phenotype Data** |
| *the percentage of VAR mice whose screen results overlap with those of B6 mice* |
| *the percentage of VAR mice whose screen results overlap with those of REF mice* |
| *difference between HET and VAR results* |
| *direction of the results (increased if the VAR average is greater than REF average)* |
| *difference between REF and VAR results* |
| *number of female HET mice* |
| *number of female REF mice* |
| *number of male REF mice* |
| *number of male HET mice* |
| *number of male VAR mice* |
| *number of female VAR mice* |
| *the identity of the phenotype (e.g. FACS T cell)* |
| *the group identity of the phenotype (e.g. FACS screen or Bone screens)* |
| *the number of outliers in REF mice* |
| *the number of outliers in HET mice* |
| *the number of outliers in VAR mice* |
| *difference between REF and B6 results* |
| *difference between REF and HET results* |
| *whether the variance of REF is big* |
| *whether the variance of HET is big* |
| *whether the variance of VAR is big* |
| *whether the average age of the mice for this mutation/phenotype is older than the average age of all mice tested this phenotype* |
| *whether the average age of the VAR mice is younger than the average age of the REF mice* |
| *number of pedigrees this gene/phenotype has* |
| *the direction of the position superpedigree results for this mutation/phenotype* |
| *how many pedigrees contributed to the significant position superpedigree results for this mutation/phenotype* |
| *number of pedigrees included in the significant position superpedigree results for this mutation/phenotype* |
| *the direction of the gene superpedigree results (null alleles) for this phenotype* |
| *the direction of the gene superpedigree results (null+missense alleles) for this phenotype* |
| *whether there are corresponding trimmed results for the untrimmed data with increased direction\** |
| *how closely VAR results resemble B6 results* |
| *how closely REF results resemble B6 results* |
| *whether REF and B6 are different* |
| **Linkage Data** |
| *average number of significant Linkage Analyzer runs for each allele of this gene* |
| *number of phenotypes with significant selective gene superpedigree results for this gene* |
| *number of significant Linkage Analyzer runs for this gene* |
| *number of pedigrees in the selective gene superpedigree and whether the result is significant for this gene/phenotype* |
| *number of pedigrees contributing to a significant gene superpedigree result (null alleles)* |
| *number of pedigrees in a significant gene superpedigree run (null alleles)* |
| *the minimum p-value of single Linkage Analyzer for this mutation/phenotype* |
| *the percentage of body weight screens that are significant* |
| *the percentage of FACS screens that are significant* |
| *whether the gene superpedigree results are significant (null+missense)* |
| whether p-value is significant in both raw and normalized assays for this mutation/phenotype |
| whether the minimum p-value is recessive |
| whether this phenotype is driven by another mutation |
| the percentage of DSS screens that are significant |
| number of significant FACS phenotypes for this mutation |
| number of significant DSS phenotypes for this mutation |
| number of significant body weight phenotypes for this mutation |
| whether the position superpedigree results are significant for this mutation/phenotype |
| whether the gene superpedigree results are significant (null alleles) |
| whether the gene superpedigree results are significant (missense alleles) for this phenotype |
| **Mutation Data** |
| *damage score for this mutation* |
| number of alleles this gene has |
| whether the mutation is autosomal |
| whether the mutation is co-localized with another mutation for this phenotype |
| whether the mutation is co-localized with a verified mutation for this phenotype |
| whether the mutation is co-localized with an excluded mutation for this phenotype |
| whether the mutation is co-localized with a mutation of higher damage score |
| the number of splice variants for this mutation |
| **Gene Data** |
| the p-value for a lethal phenotype |
| the probability that the gene is an essential gene (Essentiality Score) |
| **Other** |
| *number of phenotypes with Algorithmic Score greater or equal to -0.5 for this mutation* |
| *Algorithmic Score for this mutation/phenotype* |

\*Trimmed results = untrimmed data normalized for cell viability.
The top 20 most important features are italicized.

**Table 4.** Top 20 most important features of input data in CE prediction algorithm

|  |  |
| --- | --- |
| 1 | number of phenotypes with Algorithmic Score greater or equal to -0.5 for this mutation |
| 2 | average number of significant Linkage Analyzer runs for each allele of this gene |
| 3 | Algorithmic Score for this mutation/phenotype |
| 4 | number of phenotypes with significant selective gene superpedigree results for this gene |
| 5 | number of significant Linkage Analyzer runs for this gene |
| 6 | damage score for this mutation |
| 7 | number of pedigrees in the selective gene superpedigree and whether the result is significant for this gene/phenotype |
| 8 | number of pedigrees contributing to a significant gene superpedigree result (null alleles) |
| 9 | number of pedigrees in a significant gene superpedigree run (null alleles) |
| 10 | the percentage of VAR mice whose screen results overlap with those of B6 mice |
| 11 | the minimum p-value of single Linkage Analyzer for this mutation/phenotype |
| 12 | the percentage of body weight screens that are significant |
| 13 | the percentage of FACS screens that are significant |
| 14 | the percentage of VAR mice whose screen results overlap with those of REF mice |
| 15 | difference between HET and VAR results |
| 16 | direction of the results (increased if the VAR average is greater than REF average) |
| 17 | whether the gene superpedigree results are significant (null+missense) |
| 18 | difference between REF and VAR results |
| 19 | number of female HET mice |
| 20 | number of female REF mice |

\*Ranked from most (1) to least (20) important

#### Damage Score

The Damage Score (range 0-1), a mutation feature, is the sixth most important feature overall in the CE algorithm. The Damage Score denotes the likelihood that a protein is functionally impaired and is determined by a machine learning algorithm that integrates 38 independent prediction scores from the human database for Non Synonymous Functional Prediction (dbNSFP) and the probability of protein damage to phenovariance caused by mouse mutations^3^. A higher score suggests a mutation is more likely to be deleterious. The current Damage Score prediction model was trained on 871 known deleterious mutations and 1,797 known neutral mutations. 666 mutations with known effects were used to test the performance of the established model, which yielded a ROC curve with area under the curve (AUC) of 0.852 (Supplementary Fig. 2).

#### Essentiality Score

The Essentiality Score (range 0-1) is a gene feature and denotes the likelihood of lethality prior to weaning age (4 weeks postpartum) in mice homozygous for a robust knockout allele of a gene. The Essentiality Score is calculated using a machine learning algorithm incorporating various independent features of genes including gene conservation, protein-protein interaction network, expression stage, and viability/proliferative ability of human cell lines in which the gene is mutated. The Essentiality Score prediction model is trained at monthly intervals. The current training data set consists of 3,500 known non-essential genes (Essentiality Score = 0) and 2,058 known essential genes (Essentiality Score = 1), determined based on annotations in the Mouse Genome Informatics (MGI) Database and observed effects of CRISPR-targeted null mutations we generated in C57BL/6J mice. The current cutoff values are > 0.55 for essential genes and < 0.45 for non-essential genes, and are used to inform gene targeting efforts, in which either a knockout allele or a replacement identical to the original ENU allele is created for verification of phenotype. 1,389 genes with known effects on viability were used to test the performance of the established model, yielding a ROC curve with AUC of 0.893 (Supplementary Fig. 3).

#### Algorithmic Score

Assessments of mutation-phenotype associations are made using a human-developed algorithm that outputs a points-based score called the Algorithmic Score (current range −13.5-3.5). The Algorithmic Score appears twice among the most important features contributing to the CE algorithm (first and third in importance; Table 4). The algorithm consists of a set of rules based on empirical observations (Table 5). For each feature supporting or opposing the authenticity of a mutation-phenotype association, respectively, the Algorithmic Score is increased or decreased. The features used in the Algorithmic Score calculation are similar to those used in the CE machine learning algorithm, but static (not influenced by exposure to new training data), and the performance of the rule-based algorithm by itself falls short of the performance of the CE prediction model (Fig. 1d).

**Table 5.**
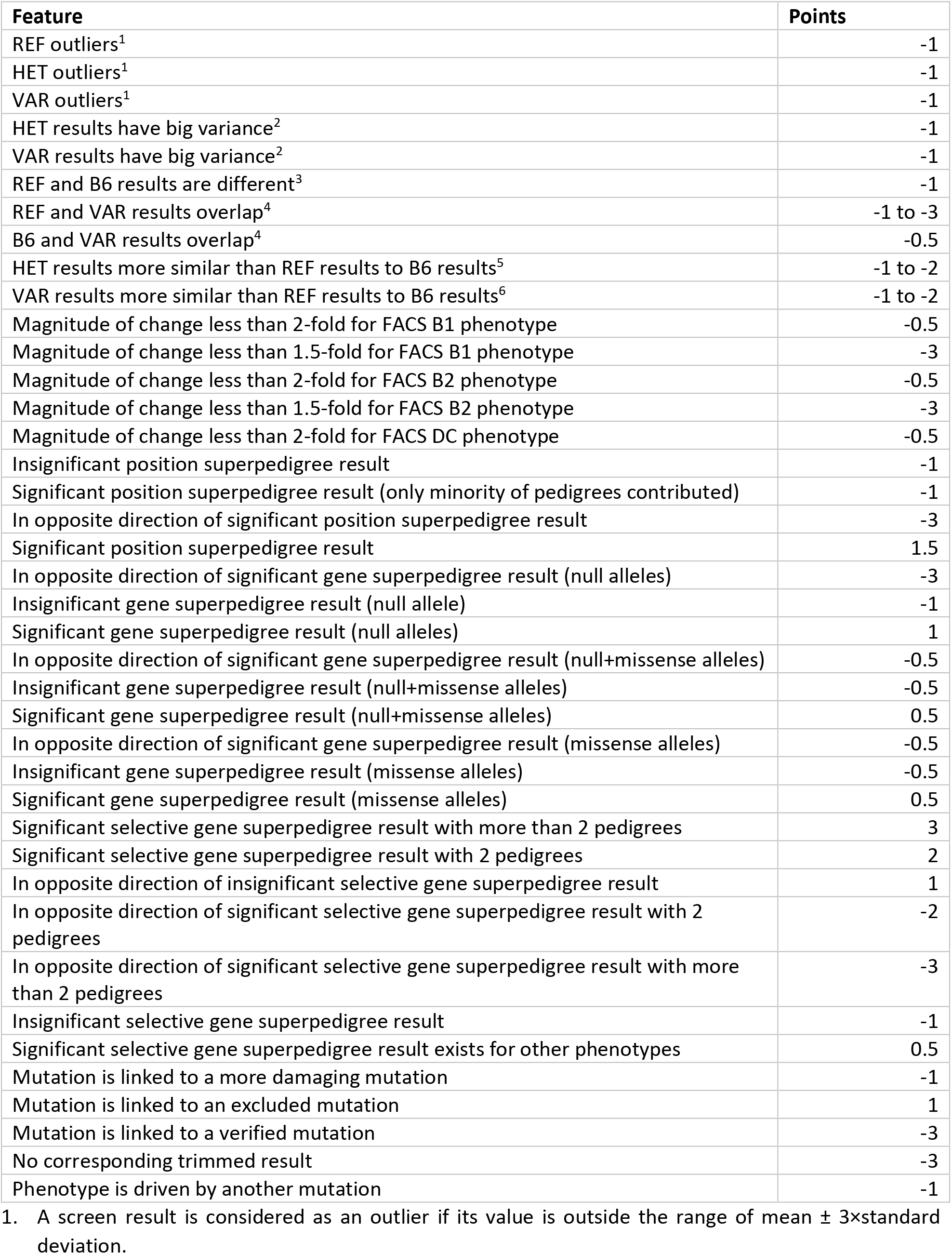
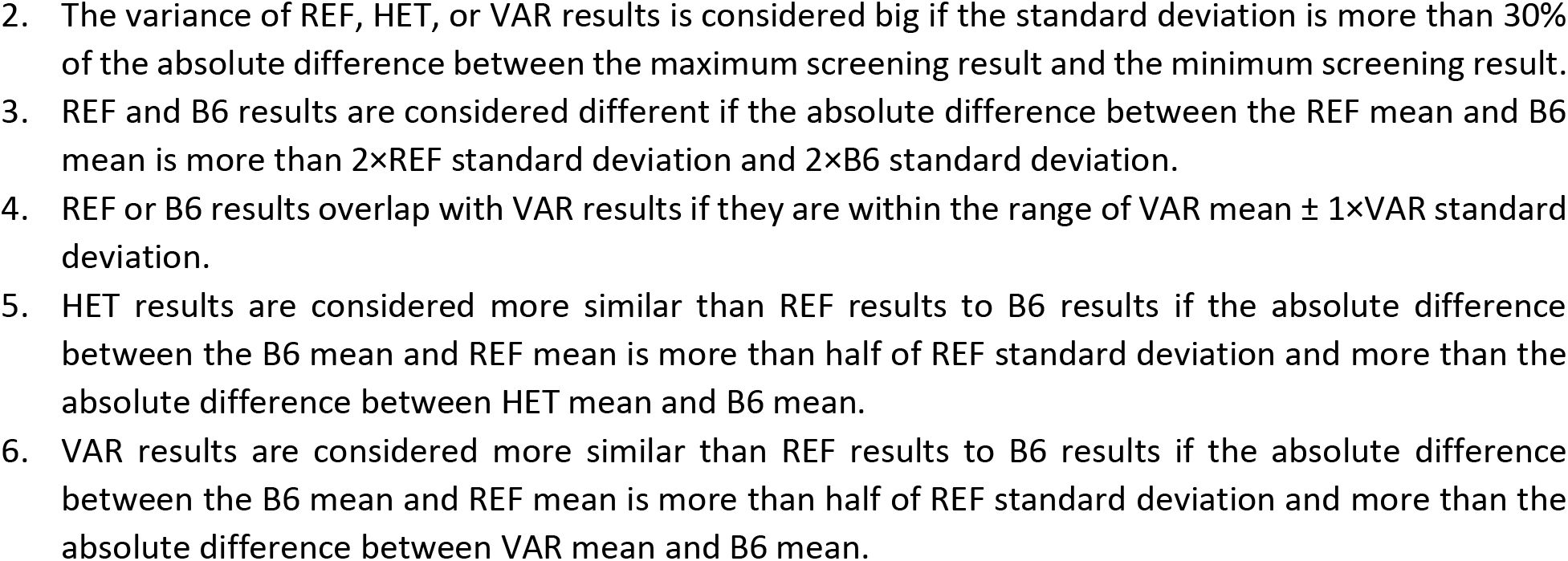
Rules for Algorithmic Score determination

| **Feature** | **Points** |
| --- | --- |
| REF outliers<sup>1</sup> | -1 |
| HET outliers<sup>1</sup> | -1 |
| VAR outliers<sup>1</sup> | -1 |
| HET results have big variance<sup>2</sup> | -1 |
| VAR results have big variance<sup>2</sup> | -1 |
| REF and B6 results are different<sup>3</sup> | -1 |
| REF and VAR results overlap<sup>4</sup> | -1 to -3 |
| B6 and VAR results overlap<sup>4</sup> | -0.5 |
| HET results more similar than REF results to B6 results<sup>5</sup> | -1 to -2 |
| VAR results more similar than REF results to B6 results<sup>6</sup> | -1 to -2 |
| Magnitude of change less than 2-fold for FACS B1 phenotype | -0.5 |
| Magnitude of change less than 1.5-fold for FACS B1 phenotype | -3 |
| Magnitude of change less than 2-fold for FACS B2 phenotype | -0.5 |
| Magnitude of change less than 1.5-fold for FACS B2 phenotype | -3 |
| Magnitude of change less than 2-fold for FACS DC phenotype | -0.5 |
| Insignificant position superpedigree result | -1 |
| Significant position superpedigree result (only minority of pedigrees contributed) | -1 |
| In opposite direction of significant position superpedigree result | -3 |
| Significant position superpedigree result | 1.5 |
| In opposite direction of significant gene superpedigree result (null alleles) | -3 |
| Insignificant gene superpedigree result (null allele) | -1 |
| Significant gene superpedigree result (null alleles) | 1 |
| In opposite direction of significant gene superpedigree result (null+missense alleles) | -0.5 |
| Insignificant gene superpedigree result (null+missense alleles) | -0.5 |
| Significant gene superpedigree result (null+missense alleles) | 0.5 |
| In opposite direction of significant gene superpedigree result (missense alleles) | -0.5 |
| Insignificant gene superpedigree result (missense alleles) | -0.5 |
| Significant gene superpedigree result (missense alleles) | 0.5 |
| Significant selective gene superpedigree result with more than 2 pedigrees | 3 |
| Significant selective gene superpedigree result with 2 pedigrees | 2 |
| In opposite direction of insignificant selective gene superpedigree result | 1 |
| In opposite direction of significant selective gene superpedigree result with 2 pedigrees | -2 |
| In opposite direction of significant selective gene superpedigree result with more than 2 pedigrees | -3 |
| Insignificant selective gene superpedigree result | -1 |
| Significant selective gene superpedigree result exists for other phenotypes | 0.5 |
| Mutation is linked to a more damaging mutation | -1 |
| Mutation is linked to an excluded mutation | 1 |
| Mutation is linked to a verified mutation | -3 |
| No corresponding trimmed result | -3 |
| Phenotype is driven by another mutation | -1 |

1. A screen result is considered as an outlier if its value is outside the range of mean ± 3×standard deviation.
2. The variance of REF, HET, or VAR results is considered big if the standard deviation is more than 30% of the absolute difference between the maximum screening result and the minimum screening result. 3. REF and B6 results are considered different if the absolute difference between the REF mean and B6 mean is more than $2 \times \text{REF standard deviation}$ and $2 \times \text{B6 standard deviation}$ . 4. REF or B6 results overlap with VAR results if they are within the range of $\text{VAR mean} \pm 1 \times \text{VAR standard deviation}$ . 5. HET results are considered more similar than REF results to B6 results if the absolute difference between the B6 mean and REF mean is more than half of REF standard deviation and more than the absolute difference between HET mean and B6 mean. 6. VAR results are considered more similar than REF results to B6 results if the absolute difference between the B6 mean and REF mean is more than half of REF standard deviation and more than the absolute difference between VAR mean and B6 mean.

#### Driven By status

Another input feature to the CE algorithm is generated by a software program called Driven By, which evaluates both linked and unlinked candidate mutations to determine the best candidate. At times a cluster of linked mutations fails to undergo meiotic separation; hence more than one mutation may stand as a candidate for causation of a phenotype. On other occasions, as a matter of happenstance, homozygotes for a non-causative, unlinked mutation may also be homozygous for a causative mutation. Usually this occurs when the number of homozygotes for the non-causative mutation is small. The Driven By program omits all instances of shared zygosity for both mutations and re-computes P values testing departure from the null hypothesis in recessive, additive, and dominant models of transmission, and determines which mutation is the more robust causation candidate. This mutation is assigned “driver” status. Based on other factors (e.g., which mutation is the most damaging, which mutation is the most essential for survival to weaning age, and which mutation has evidence of other alleles with a similar phenotype), CE may be able to identify the causative mutation out of a set of colocalizing mutations, giving it a markedly superior ML score.

Finally, an allelic series probed with a phenotypic screen provides an extremely important clue to causation and is considered in CE assessments (Tables 3 and 4, multiple rows). Superpedigrees— composites of multiple pedigrees with different alleles assayed in the same screen—are of three types. “Gene superpedigrees” pool different and identical alleles of a given gene, subjected to the same screen. “Position superpedigrees” pool identical alleles only. Identical alleles may result from 1) chance mutation of the same nucleotide; 2) transmission of a single mutation to multiple G1 descendants of a single G0 mouse; and 3) a background mutation present in mutagenized stock and shared by multiple G0 mice. “Selective gene superpedigrees” incorporate only alleles associated with *P* values ≤ 0.05 with a common direction of effect in a given phenotypic screen, and thus give an intentionally biased view of mutation effects. Because many (but not all) ENU-induced mutations are functionally hypomorphic, a selective gene superpedigree for a set of mutations in a particular gene can strongly implicate that gene in the phenotype probed by the screen in question. The number of pedigrees (and alleles) tested is also important; for very large genes, hundreds of alleles may have been tested, and the finding that two or three alleles score in a particular screen may be due to chance alone. CE takes account of this in computing probability of causation (Table 4, #7).

### CE assessments of 81,961 mutation-phenotype associations identified by AMM in flow cytometry screens

The flow cytometry screens survey 42 parameters of peripheral blood cells, measuring the frequencies of various immune cell populations and expression levels of several cell surface markers (Table 6). Of 6,680,105 mutation-phenotype associations tested by AMM in the flow cytometry screens, 82,373 passed the default initial filters, permitting analysis by CE. These putative mutation-phenotype associations emanated from 37,292 mutations in 14,437 genes, resident in 128,911 G3 mice from 3,664 pedigrees. Restriction to “good” or “excellent” candidates reduced the number of mutation-phenotype associations to 5,982, emanating from 1,768 mutations in 1,005 genes, resident in 1,297 pedigrees (Supplementary Data 1; see also CE online for the most updated data set). Gene-phenotype associations for the 1,005 genes (those with at least one good/excellent mutation-phenotype association) are displayed in a heatmap in Supplementary Data 2.

**Table 6.** Flow cytometry screening parameters

|  |  |
| --- | --- |
| 1 | B cells |
| 2 | B:T cell ratio |
| 3 | B-1 B cells |
| 4 | B-1a B cells |
| 5 | B-1a B cells in B-1 B cells |
| 6 | B-1b B cells |
| 7 | B-1b B cells in B-1 B cells |
| 8 | B-2 B cells |
| 9 | B220 MFI |
| 10 | CD11b+ DC (gated in CD11c+ cells) |
| 11 | CD11c+ DC |
| 12 | CD4:CD8 T cell ratio |
| 13 | CD4+ T cells |
| 14 | CD4+ T cells in CD3+ T cells |
| 15 | CD44 MFI on CD4+ T cells |
| 16 | CD44+ CD4+ T cells |
| 17 | CD44 MFI on CD8+ T cells |
| 18 | CD44+ CD8+ T cells |
| 19 | CD44+ T cells |
| 20 | CD44 MFI on T cells |
| 21 | CD8+ T cells |
| 22 | CD8+ T cells in CD3+ T cells |
| 23 | CD8α+ DC (gated in CD11c+ cells) |
| 24 | Central memory CD4+ T cells in CD4+ T cells |
| 25 | Central memory CD8+ T cells in CD8+ T cells |
| 26 | Effector memory CD4+ T cells in CD4+ T cells |
| 27 | Effector memory CD8+ T cells in CD8+ T cells |
| 28 | Effector T cells |
| 29 | IgD MFI |
| 30 | IgD+ B cells |
| 31 | IgM MFI |
| 32 | IgM+ B cells |
| 33 | Macrophages |
| 34 | Memory T cells |
| 35 | Naïve CD4+ T cells in CD4+ T cells |
| 36 | Naïve CD8+ T cells in CD8+ T cells |
| 37 | Naïve T cells |
| 38 | Neutrophils |
| 39 | NK cells |
| 40 | NK1.1+ T cells |
| 41 | Plasmacytoid DC |
| 42 | T cells |

Parameters represent frequencies unless otherwise indicated. MFI, mean fluorescence intensity.

We could make several observations concerning gene-phenotype associations (Supplementary Data 2). First, mutations in the majority (696 genes, 69.4%) of the 1,005 genes resulted in three or fewer good/excellent phenotype associations, with 438 genes (43.6%) having a single good/excellent phenotype association (Fig. 2a). In contrast, only 26 genes (2.6%) had at least 20 good/excellent phenotype associations, and among them 21 are well known immune regulatory genes. Second, we found that the number of good/excellent gene associations varied widely depending on the cell type affected, with B cell and T cell phenotypes associated with the most genes and conventional and plasmacytoid dendritic cell phenotypes associated with very few genes (Fig. 2b). Finally, 347 genes (34.6%) known or predicted to be essential for viability (Essentiality Score > 0.55) were associated with at least one flow cytometry phenotype, indicating that numerous developmentally important genes likely also have postnatal functions in leukocytes (Fig. 2c).

**Figure 2.**
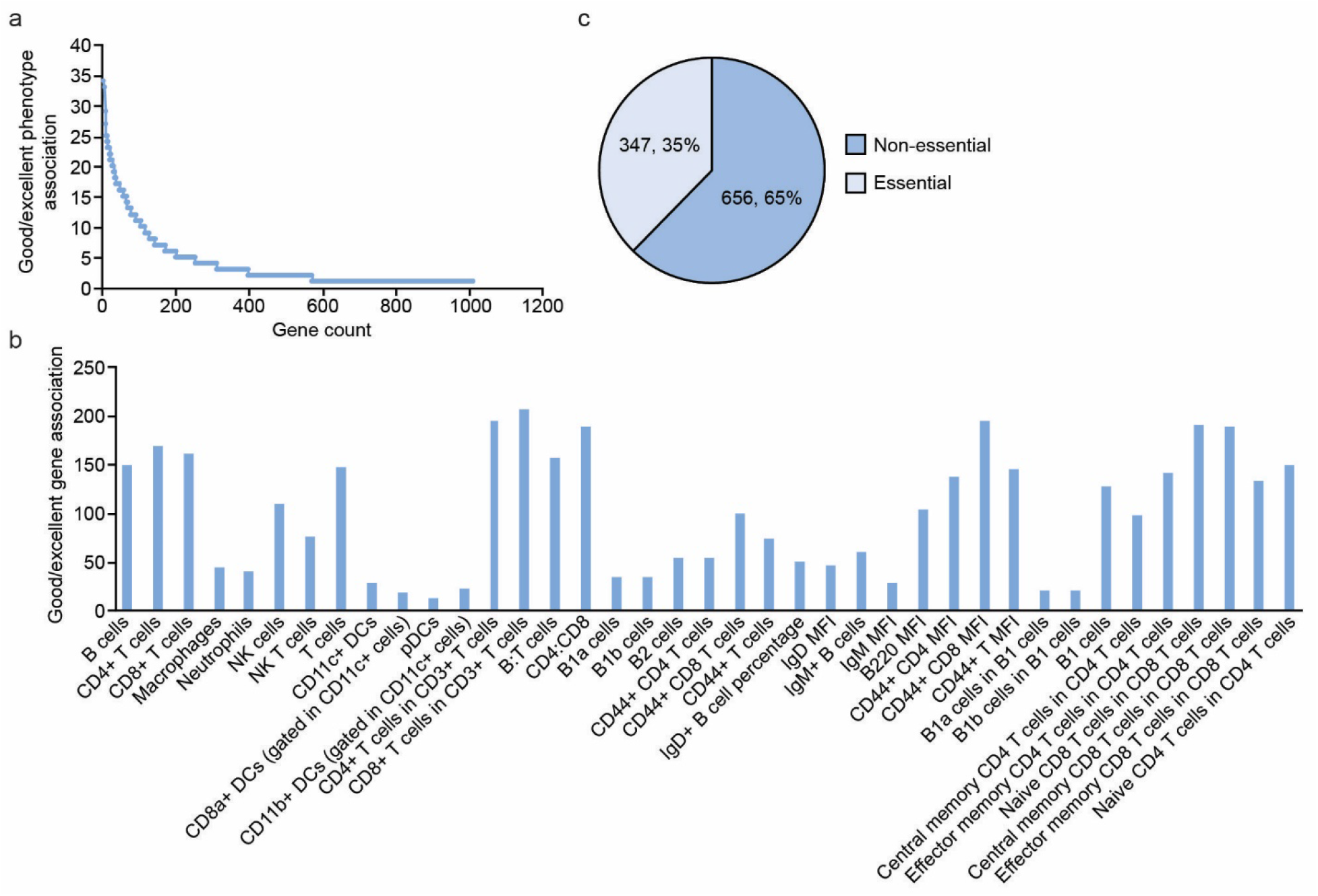
Characteristics of gene-phenotype associations for 1,005 genes with at least one good/excellent mutation-phenotype association. (**a**) Number of good/excellent phenotype associations plotted versus gene count. (**b**) Number of good/excellent gene associations plotted versus flow cytometry parameter. Parameters are cell frequencies unless indicated. MFI, mean fluorescence intensity. (**c**) Number and percentage of essential and non-essential genes.

A total of 1,045 mutations in 521 genes rated good/excellent by CE and suspected or proven causative of flow cytometry phenotypes were given allele names and annotated as phenotypic mutations in the Mutagenetix database, irrespective of present candidate status (Supplementary Data 3). While we consider that named alleles are very likely causative, we cannot be certain that un-named alleles are not also causative. Some of the un-named alleles are designated as “linked to” or “driven by” another mutation in the same pedigree. This may indicate that they are not causative, but does not always guarantee it, and in some cases, two named alleles are linked, suggesting that we have declared both mutations to be causative (even though they may co-localize). Definitive evidence for such dual causation can only be adduced by CRISPR/Cas9 targeting.

To begin to distinguish biological processes that may be particularly important for the development or maintenance of immune cell populations, we searched for highly represented Gene Ontology (GO) annotations associated with the 521 genes with named alleles (Supplementary Data 4 and 5). As expected, the biological process annotations were most highly enriched for terms related to immune system processes (180 genes, *P* = 3.99e-41), lymphocyte activation (102 genes, *P* = 9.27e-41), immune system development (104 genes, *P* = 1.23e-36), and other immune development/regulatory processes (Fig. 3a), which was consistent with our manual evaluation identifying 213 (40.9%) of the 521 genes as previously known immune regulators (Supplementary Data 4 and 5). By manual evaluation, 308 genes represented “new” immunologically important genes, each necessary for a normal flow cytometry profile. For many of these genes, mutant alleles were not previously available in mice and no primary immunological or other phenotypic data were available. This may be due in part to known or predicted lethality caused by null alleles of 116 of these 308 genes (Essentiality Score > 0.55). When we excluded genes with GO annotations encompassed by the term “immune system process” from the enrichment analysis, striking enrichment of genes associated with cellular metabolic processes (203 genes, *P* = 9.74e-12) including protein metabolism (97 genes, *P* = 0.00258) and carbohydrate metabolism (22 genes, *P* = 0.00094) was revealed (Fig. 3b and Supplementary Data 6).

**Figure 3.**
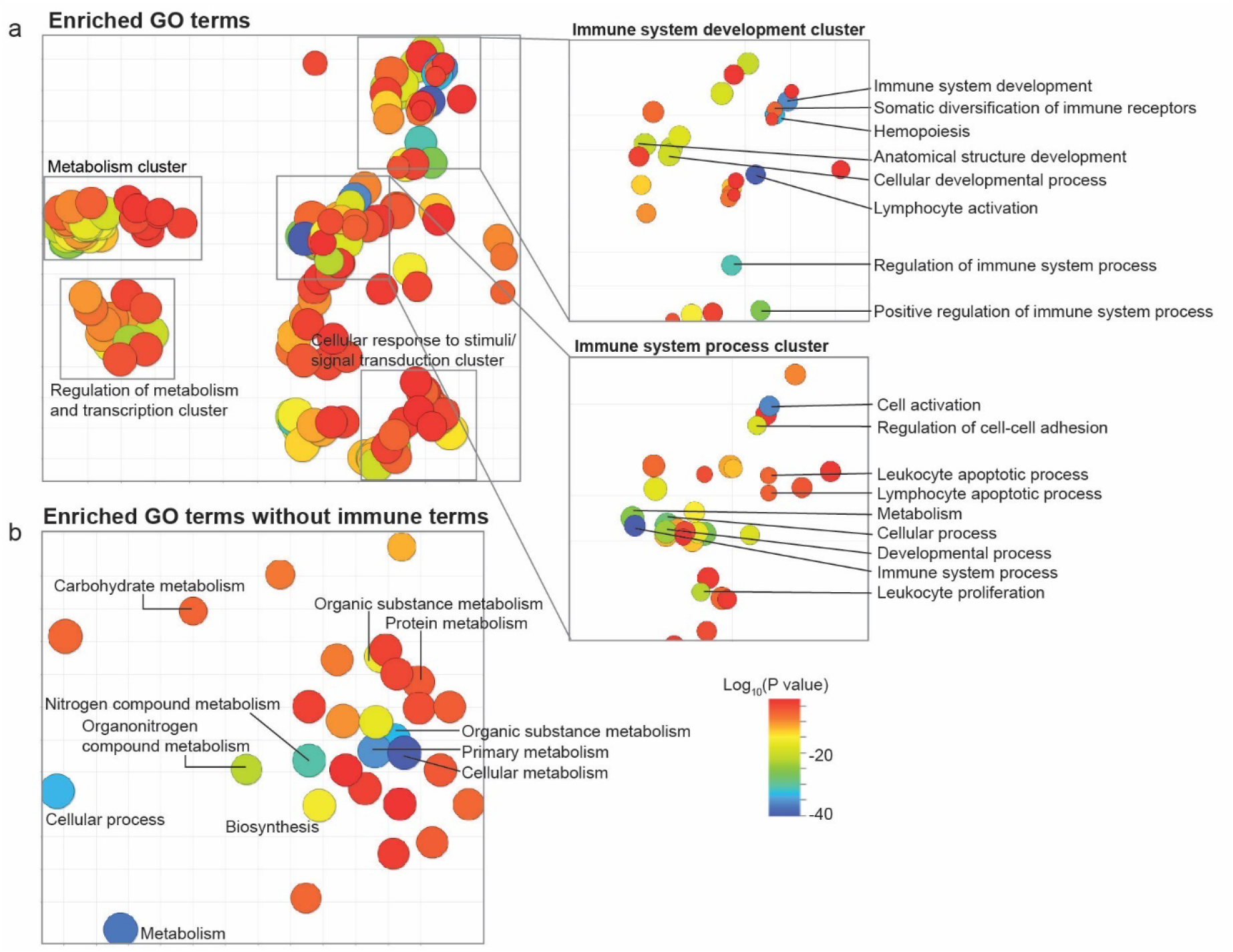
Highly represented GO annotations associated with 521 genes with named alleles. Enriched GO terms identified by GO Term Finder were distilled to a non-redundant, representative subset of the terms and displayed using REVIGO, which employs a semantic similarity-based clustering algorithm. Axes have no intrinsic meaning, but semantically similar GO terms are positioned close together so that clusters represent related biological processes. Clusters were named according to the general theme of the processes in the cluster. (**a**) GO term clusters representative of all enriched GO terms associated with 521 genes with good/excellent phenotype associations and named alleles. (**b**) Genes associated with GO term “immune system process” were removed before analysis of the same gene set as in (a). P values represent the statistical significance of the difference between the frequency of the term in the dataset and the frequency of the term in the genome.

We also performed a more inclusive analysis in which the 521 genes with named alleles were assigned to a defined set of broad GO annotations for biological processes without regard for enrichment (Supplementary Data 7). Based on its granular GO annotations, each gene was assigned to any of 71 parent GO terms to which it was related. This analysis revealed unexpected links between various types of immunodeficiency and genes involved in tissue and organ development, chromosome organization, translation, and mRNA processing, among others. Notably, among the 14 genes associated with mRNA processing, we earlier reported immune phenotypes caused by mutations in three of them (*Esrp1*^4^, *Rnps1*^5^, *Snrnp40*^6^) resulting from splicing dysfunctions. These findings suggest that human immunodeficiency diseases might sometimes originate from mutations affecting proteins needed for transcript processing and protein biosynthesis, most of which are essential for life (10/14 essential). We also noticed that 6 of the 14 genes affected 1 or 2 phenotypes only, suggesting cell type-specific effects despite the necessity of these genes for a universal cellular process.

## Discussion

CE allows rapid examination of mutations and genes strongly predicted to affect (or not to affect) phenotypes of interest measured in forward genetic screening. In general, CE is superior to the human researcher in evaluating mutation-phenotype associations because of its ability to integrate parameters not intuitively favorable or detrimental with respect to linkage analysis, and because it can perform this evaluation more rapidly on a large scale. Using the numerical ML score and categorical assessment given by CE, it is simple to rank mutations into priority lists for further in-depth study. In addition, causative mutations can frequently be discerned among several colocalizing mutations. As millions of coding/splicing mutations are introduced into the mouse genome pedigree by pedigree, more extensive allelic series will result, and nearly all genes in which causative loss-of-function mutations can exist will be identified with high confidence. CE is a tool necessary to deconvolute causation and permit this to occur.

Beyond its use as a tool for rapid identification of the mutations responsible for ENU-induced phenotypes, CE should be exceptionally useful to mouse geneticists studying complex traits (e.g., the Collaborative Cross) and to clinical geneticists concerned with the identification of rare causes of disease phenotypes. In the former case, meiotic mapping may confine phenotypes to a relatively large genomic interval, within which many candidate genes with mutational differences exist. If the phenotype is immunologic, knowledge of all genes from which flow cytometric phenotypes emanate is an important starting point for studies of causation, wherein these genes can be targeted. In the latter case, for patients with immunopathology and flow cytometric anomalies—but no mutation in a “classical” causative gene—other gene variants may be evaluated using CE. Mouse gene symbols corresponding to all loci mutated in the patient (identified by whole genome or whole exome sequencing) can be entered into CE and searched as a batch. Those found to cause a flow cytometric abnormality in the mouse evocative of that in the patient may be considered prime candidates. If genetic mapping has been performed in a human family and a particular chromosomal region has been identified, the identification of a candidate gene can be made with even higher confidence using CE, which also accepts human chromosome coordinates as search input. Moreover, a mutant mouse can, in most cases, be ordered immediately from the MMRRC, providing a model of the patient for laboratory study. Because the majority of mutations cause loss of function (rather than gain of function or new functions), and the majority of mouse genes have human orthologues or homologues, many such cases might quickly be solved.

In this paper, we evaluated mutation-phenotype associations representative of 14,437 genes with one or more variant alleles and 42 flow cytometric parameters of peripheral blood leukocytes. Flow cytometric analyses allow detection and measurement of immune cell populations with specific functional correlates, and provide insight into the developmental stages cells traverse. Abnormal flow cytometry patterns are often associated with immune dysfunction, and many immunodeficiency and autoimmune phenotypes were initially detected not by functional screens per se, but by analyzing the peripheral blood with flow cytometry. Human disease states, exhibiting similar or identical flow cytometry phenotypes, attest to the clinical relevance of many mouse flow cytometry abnormalities^7, 8, 9, 10, 11, 12, 13, 14^. We have to date achieved approximately 50% genome saturation in screening 42 flow cytometry parameters, from which we identified 777 genes with good/excellent phenotype associations not previously associated with immune function. Thus, even with a false discovery rate up to 17%, we expect that about 644 more “new” immunologically important genes remain to be found.

In broadly surveying all 1,005 genes with at least one good/excellent phenotype association, we observed that a far greater percentage of genes had one, two, or three good/excellent phenotype associations (69.3%) compared to the percentage with many (≥20) good/excellent phenotype associations (2.6%). These findings suggest that the majority of genes affecting immune cell populations in the blood carry out cell type- or phenotype-specific functions. We are investigating the hypothesis that identical or similar combinations of phenotypes affected by two or more genes can indicate the functioning of those genes in a common molecular pathway. We also observed that good/excellent gene associations did not affect cell populations with equal frequency despite uniform phenotypic testing across all screens. For example, CD4+ or CD8+ T cells had 3.6-to 13.1-fold more gene associations than plasmacytoid DC, macrophages, or neutrophils. While a trivial explanation is that significant phenotypic differences are detected less often for rarer blood cell populations, another possibility reflecting the biology of the cells is that T cells are intrinsically less tolerant of genetic variation than plasmacytoid DC, macrophages, or neutrophils, at least with regard to the numbers of these cells represented in the peripheral blood. An understanding of individual protein function and the pathways they regulate is critical to gain insight into these issues. To this end, we performed two types of GO analysis on the set of 521 genes with named alleles, either searching for enriched GO terms or binning genes by a static set of broad GO terms, and found several previously unappreciated processes that appear to be important in immune cell development or maintenance. These include carbohydrate metabolism, protein metabolism, mRNA processing, translation, and anatomical structure development. Future work should focus on fitting the mechanisms of each candidate gene into these larger pathways.

The vast majority of mice phenotyped by flow cytometry were also phenotyped in other screens, among them screens measuring responses to immunization, innate immune responses, body weight, skeletal measurements, blood pressure, heart rate, dextran sodium sulfate (DSS) sensitivity, circadian rhythms, and/or motor coordination. In the future, the data from other screens will be released for public users of CE to interpret a wide range of phenotypic consequences that emanate from each mutation. All biomedically relevant phenotypic screens may ultimately enlighten the study of human phenotype and help to distinguish mechanisms of phenotypes caused by certain alleles, as many mutations score in disparate screens (for example, immune function and body weight, or immune function and neurobehavioral function).

## Methods

### Mice

Eight-to ten-week old C57BL/6J males purchased from The Jackson Laboratory were mutagenized with ENU as described previously^15^. Mutagenized G0 males were bred to C57BL/6J females, and the resulting G1 males were crossed to C57BL/6J females to produce G2 mice. G2 females were backcrossed to their G1 sires to yield G3 mice, which were screened for phenotypes. Whole-exome sequencing and mapping were performed as described^1^.

To generate mice carrying CRISPR/Cas9-targeted mutations, female C57BL/6J mice were superovulated by injection with 6.5 U pregnant mare serum gonadotropin (PMSG; Millipore), then 6.5 U human chorionic gonadotropin (hCG; Sigma-Aldrich) 48 hours later. The superovulated mice were subsequently mated with C57BL/6J male mice overnight. The following day, fertilized eggs were collected from the oviducts and *in vitro* transcribed Cas9 mRNA (50 ng/μl) and small base-pairing guide RNA (50 ng/μl) were injected into the cytoplasm or pronucleus of the embryos. The injected embryos were cultured in M16 medium (Sigma-Aldrich) at 37°C and 5% CO_2_. For the production of mutant mice, 2-cell stage embryos were transferred into the ampulla of the oviduct (10-20 embryos per oviduct) of pseudo-pregnant Hsd:ICR (CD-1) (Harlan Laboratories) females.

Mice were housed in specific pathogen-free conditions at the University of Texas Southwestern Medical Center and all experimental procedures were performed in accordance with the guidelines established by the Institutional Animal Care and Use Committee of the University of Texas Southwestern Medical Center and with the National Institutes of Health Guide for the Care and Use of Laboratory Animals. Male and female mice were used in all experiments and data for males and females were combined for analysis.

### FACS

Peripheral blood was collected from G3 mice >6 weeks old by cheek bleeding. Red blood cells (RBCs) were lysed with hypotonic buffer (eBioscience). Samples were washed with FACS staining buffer (PBS with 1% (w/v) BSA) one time and then centrifuged at 500 × g for 5 minutes. The RBC-depleted samples were stained for 1 hour at 4°C, in 100 μl of a 1:200 cocktail of fluorescence-conjugated antibodies to 15 cell surface markers encompassing the major immune lineages B220 (BD, clone RA3-6B2), CD19 (BD, clone 1D3), IgM (BD, clone R6-60.2), IgD (Biolegend, clone 11-26c.2a), CD3ε (BD, clone 145-2C11), CD4 (BD, clone RM4-5), CD8α (Biolegend, clone 53-6.7), CD11b (Biolegend, clone M1/70), CD11c (BD, clone HL3), F4/80 (Tonbo, clone BM8.1), CD44 (BD, clone 1M7), CD62L (Tonbo, clone MEL-14), CD5 (BD, clone 53-7.3), CD43 (BD, clone S7), NK 1.1 (Biolegend, clone OK136)) and 1:200 Fc block (Tonbo, clone 2.4G2). Flow cytometry data were collected on a BD LSR Fortessa and the proportions of immune cell populations in each G3 mouse were analyzed with FlowJo software. The resulting phenotypic data were uploaded to Mutagenetix for automated mapping of causative alleles.

### Automated meiotic mapping

AMM was performed as previously described^1^. Briefly, genotypes at all mutation sites present in the exomes of G3 mice were determined prior to phenotypic screening: tail DNA from G1 males was subjected to whole exome sequencing using an Illumina HiSeq 2500 instrument; G2 and G3 mice were then genotyped at the identified mutation sites using an Ion PGM (Life Technologies). Following phenotypic screening, linkage analysis using recessive, additive, and dominant models of inheritance was performed for every mutation in the pedigree using the program Linkage Analyzer; phenotypic data scatter plots and Manhattan plots were displayed using the program Linkage Explorer. The *P* values of association between genotype and phenotype were calculated using a likelihood ratio test from a generalized linear model or generalized linear mixed effect model and Bonferroni correction applied.

### Candidate Explorer

CE is publicly accessible at https://mutagenetix.utsouthwestern.edu/linksplorer/candidate.cfm. Linkage data obtained through screening will be released in phases according to phenotype. Blood cell flow cytometry screening data are currently available for search using CE and new data will be released as they are acquired after a six-month delay from the date of screening.

### Damage Score

The Damage Score is an ensemble score that uses a logistic regression model to integrate 38 independent prediction scores from the human <u>d</u>ata<u>b</u>ase for <u>N</u>on <u>S</u>ynonymous <u>F</u>unctional <u>P</u>rediction (dbNSFP) and the probability of damage by mouse mutations sufficient to cause phenotypic change^3^. It can be used as a quantitative prediction score to measure the likelihood of a mouse mutation being deleterious.

Our assumption is that if the mouse missense mutation is the same as the human mutation (both nucleotide and amino-acid changes), then the mutation effect in human and mouse should be similar. We therefore use human scores to predict likelihood of damage in mice.

A set of mouse ENU mutations with class tags (known damaging or neutral) was retrieved from the Mutagenetix database. The known mutation class tags come from four sources: 1) physically isolated mutations (out of linkage with all other coding/splicing mutations in the pedigree) that fall within essential genes yet can be transmitted from heterozygous G2 females and their heterozygous G1 sire to homozygous G3 mice at a ratio that does not significantly depart from Mendelian expectation, are considered neutral. 2) Conversely, isolated mutations in essential genes that are NOT transmitted to homozygosity, to the extent that homozygotes are observed at frequencies significantly beneath the expected Mendelian ratio, are considered damaging. 3) Mutations that cause qualitative (usually visible) phenotypes are considered damaging. 4) Mutations that have been verified to be significant in phenotypic screening of CRISPR replacement alleles are also considered to be damaging.

The mutations tagged as damaging or neutral were lifted-over from mouse genome to human genome (translated to the equivalent amino acid) and kept for mutations that lead to the same nucleotide and amino-acid changes in both genomes. Then we searched for corresponding human mutations in the dbNSFP database to obtain scores for all available prediction methods. The retrieved scores, combined with the probability of phenotypically detectable damage by the mutations in mice, were integrated with the input data set to fit a logistic regression model. The fitting process was implemented by the 10-fold cross-validation method using R package (caret). The constructed model (classifier) was then used to compute the score of a set of mutations with unknown class membership. The data set used for prediction was created in the same way as data set used for modeling. The score predicted by the model represents the probability of a mutation being in the damaging class. The higher the score, the more likely to be deleterious the mutation.

The input data set contains 3,334 mouse mutations, of which 1,088 are deleterious and 2,246 are neutral. In order to evaluate the performance of the constructed model in predicting the membership of the new mutation category, the input data set was randomly divided into two sets: one set consisting of 2,668 mutations (80% of original dataset, 871 deleterious mutations and 1,797 neutral mutations) was used to train and validate the logistic regression model, and a second set of the remaining 666 mutations was used to test the performance of the established model.

Quartile based correspondence between raw Damage Scores and probability of protein damage to phenovariance is shown in Supplementary Table 1.

### Essentiality Score

Essentiality score (E-score) is used to estimate the likelihood of lethality in mice when the gene is knocked out. Our approach is based on the assumption that essential and non-essential genes in mice can be distinguished by various independent features of genes. The logistic regression method is used to fit the features of known essential and non-essential genes in mice to obtain a trained model for predicting the unknown essentiality of genes.

The model uses four main categories of gene features: 1) from literature: 7 gene features, such as gene conservation, protein-protein interaction network, expression stage and etc., have been suggested to be associated with gene essentiality of many species, including mouse. 2) The essentiality of human orthologous genes: the genes required for cell proliferation and viability in tested cell lines are defined as essential genes under specific conditions. Frequency of being essential in tested human cell lines was used as a feature in our model. 3) pLI score from the ExAC (<u>p</u>robability of <u>L</u>oss-of-function <u>I</u>ntolerance): the closer the score is to 1, the more likely the gene is essential to human survival. 4) Minimum P values for an ENU targeted mouse gene obtained from the lethal model by the Linkage Analyzer program.

The phenotypic description of the 8,032 genes in MGI which may be knocked out in mice was carefully reviewed and a set of genes designated as “essential” or “non-essential” were manually curated according to the following criterion: 1) If the homozygous knockout allele is explicitly described as causing embryonic lethality, neonatal lethality, prenatal lethality, perinatal lethality or pre-weaning lethality, the gene was considered to be required to survive before weaning and was classified as an essential gene. An E-score of 1 was assigned to the gene. 2) If homozygous knockout alleles are compatible with viability, normal growth, no obvious phenotype, or some phenotype, but not apparent effect on viability, then it was classified as a non-essential gene. An E-score of 0 was assigned to the gene. In addition, an E-score of 1 was assigned to those genes verified in our CRISPR KO experiments as causing significant lethality before weaning; an E-score of 0 was assigned to genes verified in our CRISPR KO experiments as resulting in normal Mendelian ratios in crosses of heterozygous mutants.

A set of 6,947 genes, in which 2,572 were labeled as essential genes and 4,375 as non-essential genes, was integrated with the above-mentioned gene features. The resulting data set was used to fit a logistic regression model with 10-fold cross-validation. The constructed model was then used to predict the essentiality of remaining mouse genes. The predicted score is between 0 and 1. The closer the score is to 1, the more likely the gene is essential.

To assess the performance of constructed model in predicting unknown essentiality of genes, the data set used to construct the model was randomly divided into two sets: one set consisting of 5,558 genes (80% of original dataset, 3,500 non-essential genes and 2,058 essential genes) were used to train and validate the logistic regression model, and the remaining 1,389 genes were used to test the performance of the established model in the training data set.

### Algorithmic Score

Each mutation-phenotype association starts with an Algorithmic Score of zero that is adjusted according to the rules in Table 5.

### Gene Ontology analysis

Summaries of GO annotations in Supplementary Data 4 were generated using the Alliance of Genome Resources SimpleMine tool (http://tazendra.caltech.edu/~azurebrd/cgi-bin/forms/agr_simplemine.cgi). Enriched GO annotations associated with various gene lists were determined using GO TermFinder^16^ (https://go.princeton.edu/cgi-bin/GOTermFinder) set to use the *Mus musculus* annotations (MGI) and exclude evidence code “IEA” (inferred from electronic annotation). The output from GO TermFinder was processed using REVIGO to summarize and visualize enriched GO categories^17^ (http://revigo.irb.hr/). REVIGO settings: allowed similarity Medium (0.7), *Mus musculus* GO database, and semantic similarity SlimRel. GO TermMapper^16^ was used to assign genes to 71 static GO parent annotations^18^ (https://go.princeton.edu/cgi-bin/GOTermMapper).

### Data Availability

CE is publicly accessible at https://mutagenetix.utsouthwestern.edu/linksplorer/candidate.cfm. Sequences of small base-pairing guide RNA used for CRISPR/Cas9 targeting are available by request from the corresponding author.

## Acknowledgements

This work was supported by National Institutes of Health grants R01 AI125581 and U19 AI100627 (to B.B.). We thank Diantha La Vine for expert help with illustrations and the video.

## Author Contributions

Conceptualization: DX, SL, CHB, BB; Methodology: DX, SL, CHB, SH; Software: DX, SL, CHB, SH; Validation: DX, SL, CHB, SH, JHC, XZ, AL, EET, ZZ, EN-G, HS, YW, DZ, TY, JSoRelle, TM, LSun, JW, RF, AS, SS, NS, HC, GC, BH, SM, DM, BN, ER, AW, MTang, XL, PA, KK, LScott, JQ, SC, BQ, JC, RS, MTadesse, QS, JSantoyo, AB, AJ; Formal Analysis: DX, SL, CHB, SH; Investigation: JHC, XZ, AL, EET, ZZ, EN-G, HS, YW, DZ, TY, JSoRelle, TM, LSun, JW, RF, AS, SS, NS, HC, GC, BH, SM, DM, BN, ER, AW, MTang, XL, PA, KK, LScott, JQ, SC, BQ, JC, RS, MTadesse, QS, JSantoyo, AB, AJ; Data Curation: DX, SL, CHB, SH; Writing-Original Draft: EMYM, BB; Writing-Review and Editing: DX, EMYM, BB; Visualization: DX, EMYM, BB; Supervision: BB; Funding Acquisition: BB.

