## Supplementary Figures and Table for "Candidate Explorer: a tool for discovery, evaluation, and display of mutations causing significant immune phenotypes"

Takuma Misawa, RIKEN Center for the Integrative Medical Science, Laboratory for Immune Cell System, 1-7-22 Suehiro-cho, Tsurumi-ku, Yokohama City, Kanagawa, 230-0045, Japan

Lei Sun, Institute of Developmental Biology and Molecular Medicine, Fudan University, Shanghai 200433, China

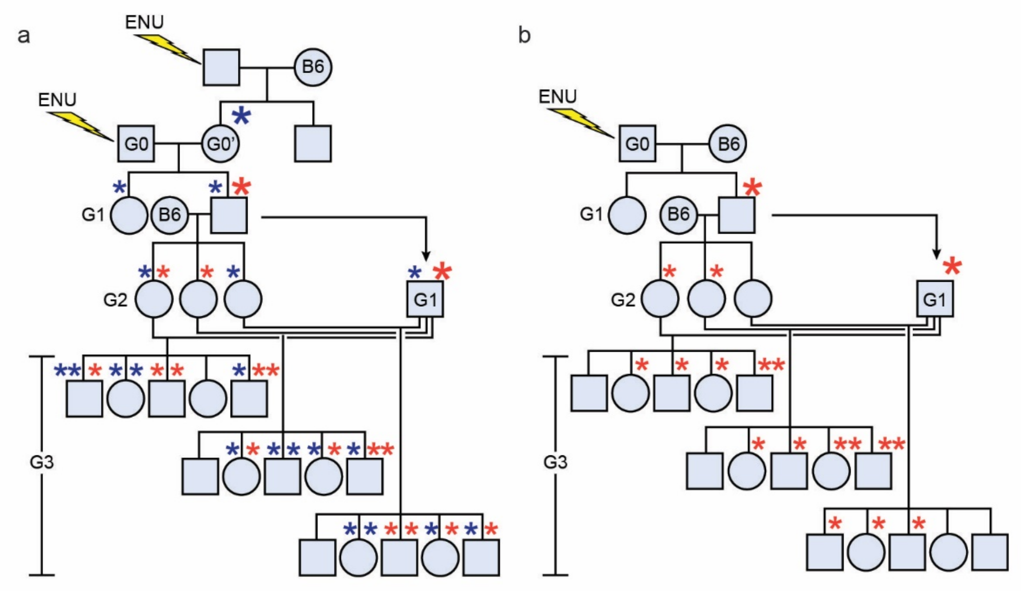


Supplementary Figure 1. Inbreeding schemes for generating G3 mice. (**a**) Mutagenized G0 males are bred to G0’ females carrying germline mutations derived from other mutagenized males. (**b**) Mutagenized G0 males are bred to wild-type C57BL/6J (B6) females. G1 males are crossed to B6 females to produce G2 mice. G2 females are crossed to their G1 father to produce G3 mice. Asterisks represent mutations derived from the G0 male (red) and G0’ female (blue); larger asterisks indicate initial germline transmission of the mutation.

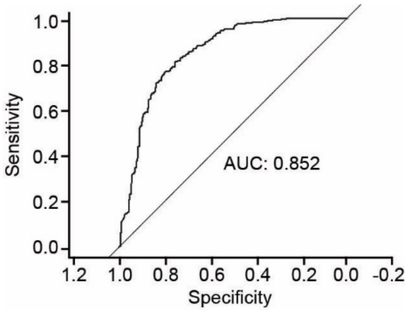


Supplementary Figure 2. ROC curve for Damage Score prediction model. Various Damage Score thresholds for calling damaging mutations were used to compute sensitivity and specificity.

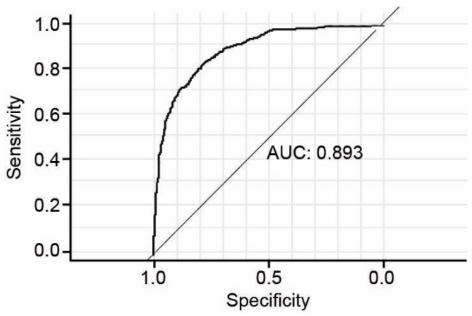


Supplementary Figure 3. ROC curve for Essentiality Score prediction model.

**Supplementary Table 1. Damage Score and probability of protein damage**

| Quartile | Damage Score | Probability (Range) of damage |
|---|---|---|
| 1 | 0.311-1.0 | 0.362 (0.27-0.452) |
| 2 | 0.184-0.311 | 0.164 (0.074-0.251) |
| 3 | 0.1-0.184 | 0.06 (0-0.145) |
| 4 | 0-0.1 | 0.044 (0-0.131) |
